# Spinal V3 interneurons and left-right coordination in mammalian locomotion

**DOI:** 10.1101/679654

**Authors:** Simon M. Danner, Han Zhang, Natalia A. Shevtsova, Joanna Borowska, Ilya A. Rybak, Ying Zhang

**Affiliations:** Department of Neurobiology and Anatomy, College of Medicine, Drexel University, Philadelphia, Pennsylvania, USA; Department of Medical Neuroscience, Brain Repair Centre, Faculty of Medicine, Dalhousie University, Halifax, Nova Scotia, Canada

**Keywords:** spinal cord, central pattern generator, locomotion, commissural neurons, V3, optogenetic stimulation, computational modeling

## Abstract

Commissural interneurons (CINs) mediate interactions between rhythm-generating locomotor circuits located on each side of the spinal cord and are necessary for left-right limb coordination during locomotion. While glutamatergic V3 CINs have been implicated in left-right coordination, their functional connectivity remains elusive. Here, we addressed this issue by combining experimental and modeling approaches. We employed *Sim1Cre/*+*; Ai32* mice, in which light-activated Channelrhodopsin-2 was selectively expressed in V3 interneurons. Fictive locomotor activity was evoked by NMDA and 5-HT in the isolated neonatal lumbar spinal cord. Flexor and extensor activities were recorded from left and right L2 and L5 ventral roots, respectively. Bilateral photoactivation of V3 interneurons increased the duration of extensor bursts resulting in a slowed down on-going rhythm. At high light intensities, extensor activity could become sustained. When light stimulation was shifted toward one side of the cord, the duration of extensor bursts still increased on both sides, but these changes were more pronounced on the contralateral side than on the ipsilateral side. Additional bursts appeared on the ipsilateral side not seen on the contralateral side. Further increase of the stimulation could suppress the contralateral oscillations by switching to a sustained extensor activity, while the ipsilateral rhythmic activity remained. To delineate the function of V3 interneurons and their connectivity, we developed a computational model of the spinal circuits consisting of two (left and right) rhythm generators (RGs) interacting via V0_V_, V0_D_ and V3 CINs. Both types of V0 CINs provided mutual inhibition between the left and right flexor RG centers and promoted left-right alternation. V3 CINs mediated mutual excitation between the left and right extensor RG centers. These interactions allowed the model to reproduce our current experimental data, while being consistent with previous data concerning the role of V0_V_ and V0_D_ CINs in securing left-right alternation and the changes in left-right coordination following their selective removal. We suggest that V3 CINs provide mutual excitation between the spinal neurons involved in the control of left and right extensor activity, which may promote left-right synchronization during locomotion.

## 1 Introduction

The rhythmic activities controlling locomotor movements in mammals are generated by neural circuits within the spinal cord, representing so-called central patter generators (CPGs; Graham Brown, 1911; Grillner, 1981, 2006; Kiehn, 2006, 2011; Rossignol et al., 2006). It is commonly accepted that each limb is controlled by a separate spinal CPG and that CPGs controlling fore and hind limbs are located in the left and right sides of lumbar and cervical enlargements, respectively. These CPGs are connected by spinal commissural interneurons (CINs) that coordinate their activities, hence defining locomotor gaits. CINs project axons across the spinal cord midline and affect interneurons and motoneurons located on the contralateral side of the cord (Butt and Kiehn, 2003; Jankowska, 2008; Kjaerulff and Kiehn, 1997; Quinlan and Kiehn, 2007).

Several classes of spinal CINs, including two subtypes of V0 CINs (excitatory V0_V_ and inhibitory V0_D_) and the excitatory V3 CIN, have been identified based on their transcription factor profiles (Gosgnach, 2011; Goulding, 2009; Lanuza et al., 2004). Both subtypes of V0 CINs promote left-right alternation, and their functional ablation resulted in aberrant left-right synchronization. *In vitro*, selective ablation of V0_D_ CINs disrupted left-right alternation at low locomotor frequencies while ablation of V0_V_ CINs disrupted left-right alternation at higher locomotor frequencies (Talpalar et al., 2013). Thus, V0 CINs are necessary for securing left-right alternation.

*In vivo*, an increase of locomotor speed in rodents is accompanied by a transition from left-right alternating gaits (walk and trot) to left-right synchronized gaits (like gallop and bound). Yet, the CIN networks promoting left-right synchronization at higher locomotor speeds, or during V0 ablation, are poorly understood. Previous modeling studies suggested that left-right synchronization could be performed by V3 CINs providing mutual excitation between the left and right rhythm-generating circuits (Danner et al., 2016, 2017; Rybak et al., 2013, 2015; Shevtsova et al., 2015; Shevtsova and Rybak, 2016). Although this suggestion allowed the previous models to reproduce the results of the above experimental studies, including the effects of V0 CIN ablation *in vitro* and *in vivo*, the role and connectivity of V3 neurons suggested by these models have not been tested experimentally and remained hypothetical.

V3 interneurons, defined by their post-mitotic expression of the transcription factor, single-minded homologue 1 (Sim1), are excitatory and the majority of them project to the contralateral side of the spinal cord (Zhang et al., 2008). Genetic deletion of V3 interneurons did not affect left-right alternation, but caused unstable gaits in walking mice, and generated imbalanced and less robust rhythmic fictive locomotion in isolated neonatal spinal cords (Zhang et al., 2008). While these experimental data strongly suggested that V3 interneurons are involved in the control of locomotion, their exact function and commissural connectivity remain mainly unknown.

To address V3’s functional connectivity between left-right spinal circuits, we took advantage of an optogenetic approach, which enabled us to specifically regulate the activity of V3 interneurons on each side of the isolated spinal cord during fictive locomotion. We then designed an updated computational model of spinal circuits that incorporated the connectivity of V3 CINs suggested from our experimental studies. Together our experimental and modeling results provide convincing evidence that V3 interneurons contribute to synchronization of the left-right locomotor activity (under appropriate conditions) by providing mutual excitation between the extensor centers of the left and right CPGs.

## 2 Results

### 2.1 Optical activation of lumbar V3 interneurons increases the intensity of extensor motor activity and slows oscillation frequency of drug-evoked fictive locomotion

To assess the function of V3 interneurons in the spinal locomotor network, we used an optogenetic approach that allowed us to selectively activate V3 interneurons in different regions of the isolated spinal cords from *Sim1Cre/*+*; Ai32* mice, which express channelrhodopsin2 (ChR2) and enhanced yellow fluorescent protein (EYFP) in Sim1 positive cells.

Using whole-cell patch-clamp recordings, we confirmed that the blue fluorescent light (450 - 490 nm) could produce membrane depolarization and evoke persistent spiking only in EYFP expressing cells from the slices of *Sim1Cre/*+*; Ai32* (Figure 1). The evoked spiking activity continued within a 20-s period with or without glutamatergic receptor blockers (CNQX and AP-5; Figure 1B,C). These results confirmed that V3 interneurons in the isolated spinal cords of *Sim1Cre/*+*; Ai32* mouse could be selectively activated by the blue fluorescent light.

**Figure 1.**
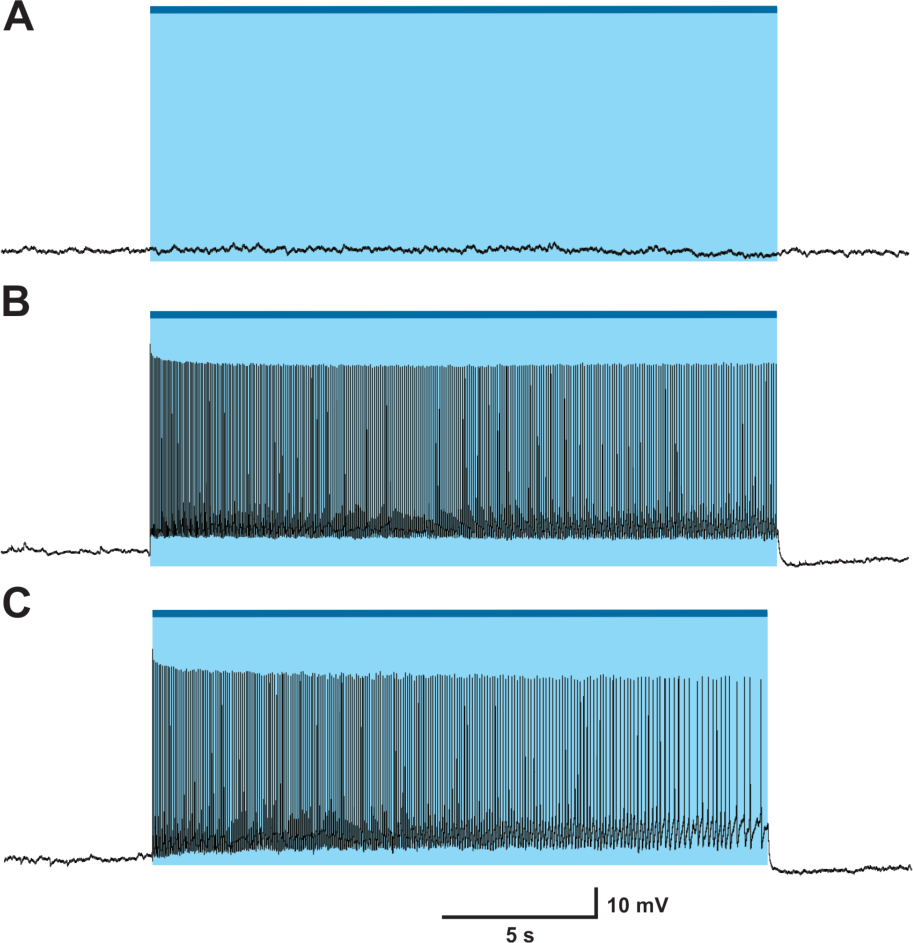
Blue LED fluorescent light evokes action potential on Sim1 cell in *Sim1*^*Cre/*+^*; Ai32* mouse. (A) Light did not activate EYFP negative cell. (B) Patch clamp recordings of *Sim1*^*Cre/*+^*; Ai32* cell with optical activation. (C) patch clamp recording of *Sim1*^*Cre/*+^*; Ai32* cell with optical activation with fast synaptic transmission blocker. Blue bar and shaded blue area represent the optical activation (light-on) period.

To investigate the role of V3 interneurons and their interactions with CPG circuits, we shone fluorescent light onto the ventral spinal cord of neonatal *Sim1Cre/*+*; Ai32* mice during fictive locomotion evoked by a 5-HT/NMDA mixture (5-HT 8 μM, NMDA 7-8 μM; Figure 2) and analyzed the effects of light stimulation on the ongoing rhythmic activity (Figure 3). Flexor and extensor activities on each side of the cord were evaluated based on the recordings from L2 and L5 ventral roots, respectively. Optogenetic stimulation with blue fluorescent light centered on the midline of the L2 ventral spinal cord (Figure 2A) slowed down the ongoing rhythmic activity, as evident by an increased locomotor cycle period of L5 ventral roots (Figures 2B and 3C; P < 0.0001). The increased cycle period was mainly attributed to an increase in the L5-burst durations, while L2-burst durations did not significantly change in response to the applied stimulation (Figures 2B and 3B). Furthermore, the optogenetic stimulation caused an increase in the amplitude of integral ENG bursts in L5, but not in L2 ventral roots (Figures 2B and 3A). These results suggest that lumbar V3 neurons interact with locomotor CPGs and provide activation of extensor circuits, either directly or transynaptically.

**Figure 2.**
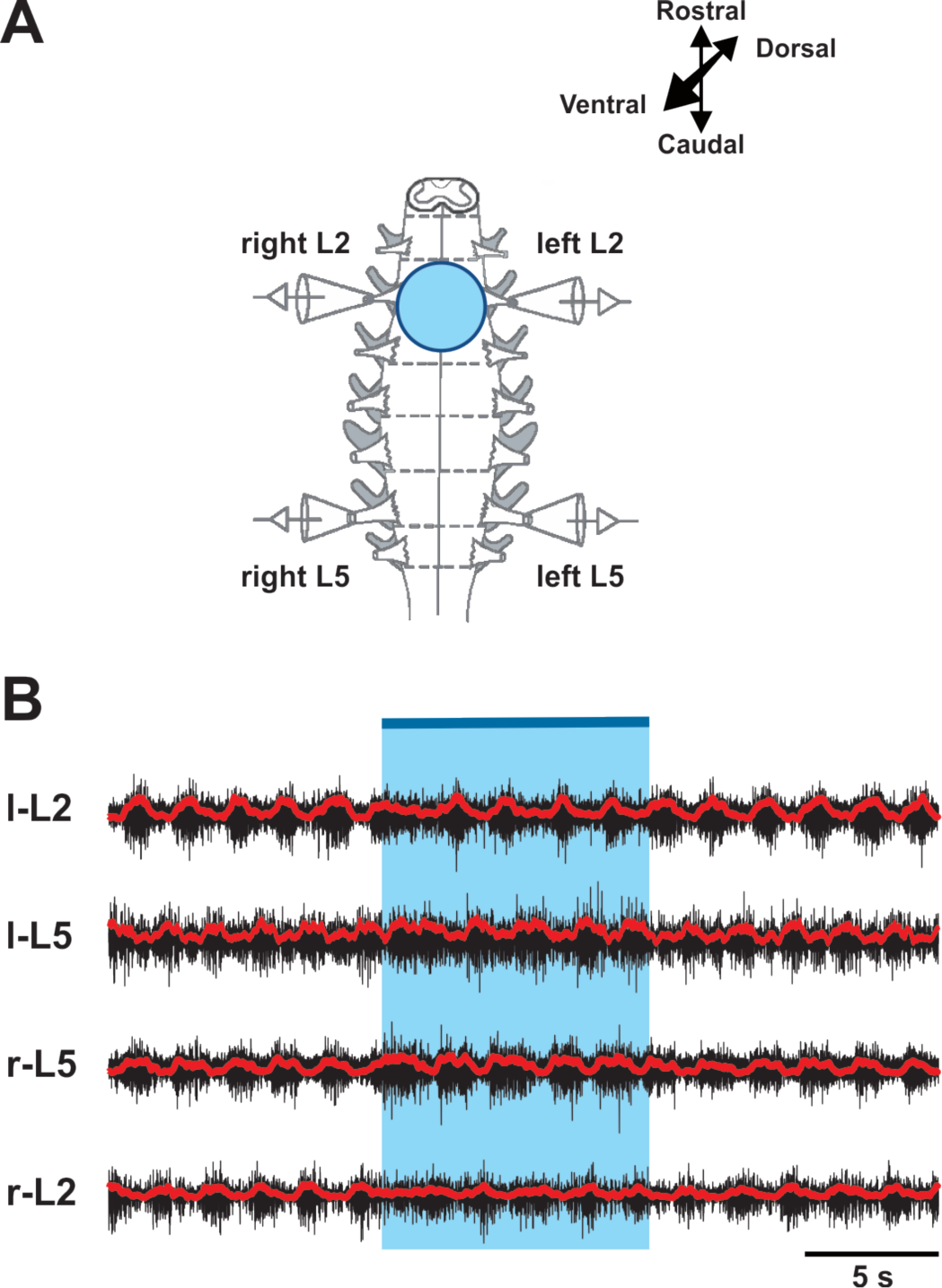
Optical activation of both sides of V3 interneurons in L2 ventral region during drug-evoked fictive locomotion. (A) Experimental setup. (B) Optical activation of lumbar V3 interneurons increases the intensity of extensor motor output and slows oscillation frequency of drug-evoked fictive locomotion. Fictive locomotion was induced by drug cocktail (5-HT 8μM, NMDA 8 μM) in *Sim1Cre/*+*; Ai32* cord. The blue bar on the top and shaded blue area represent the optical stimulation period. Red lines are rectified and smoothed traces of the original ENG recordings (black).

**Figure 3.**
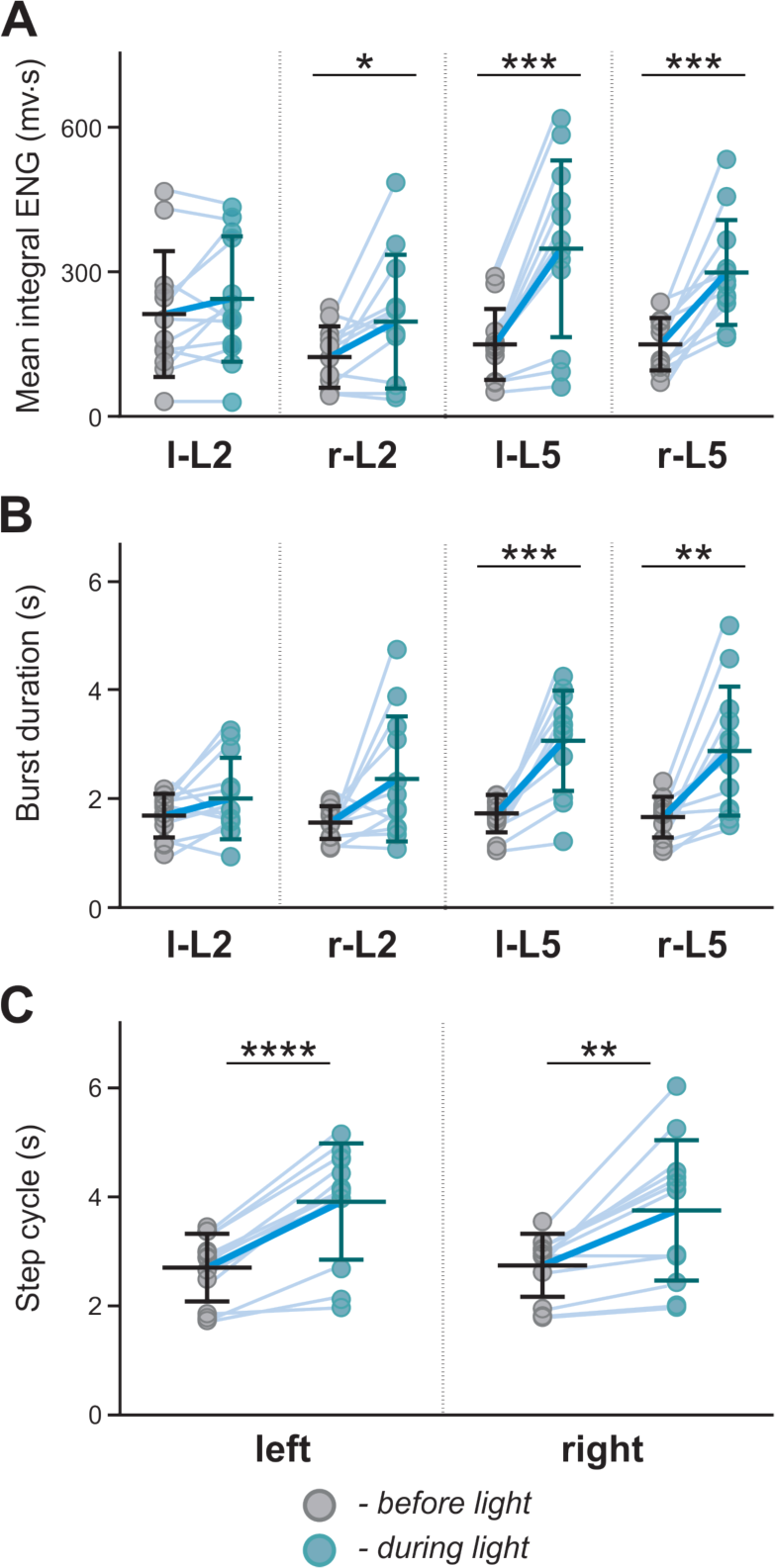
Changes in ENG characteristics by symmetric optical stimulation of V3 interneurons in spinal segment L2. Mean integral ENG (A), burst duration (B) and L5 step cycle duration (C) during drug-induced locomotor-like activity (n = 12) before (gray dots) and during (blue dots) optical activation. Each dot represents the average value of one trace. Light blue lines connecting two dots represent increase in one cord. Thick, dark blue line represent the average of increase. * indicates P < 0.05, ** indicates 0.01 < P <0.05, *** indicates 0.0001 < P <0.001, **** indicates P <0.0001.

### 2.2 Biased optical activation of V3 interneurons in spinal segment L2 leads to asymmetrical left-right motor activity

Since most of V3s are commissural interneurons, activation of V3s on one side of the spinal cord should more strongly impact the contralateral circuits. To test this hypothesis, we used a 20x, 1.0 numerical aperture (NA) objective to deliver the light onto a small area in one side of the spinal cord (Figure 4A). We then selected the illuminated region by manually adjusting the field diaphragm to have the activation zone between approximately one-third to a half of the spinal cord (Figure 4B) and the intensity of the light-emitting diode (LED) light to regulate the number of V3 neurons being activated.

**Figure 4.**
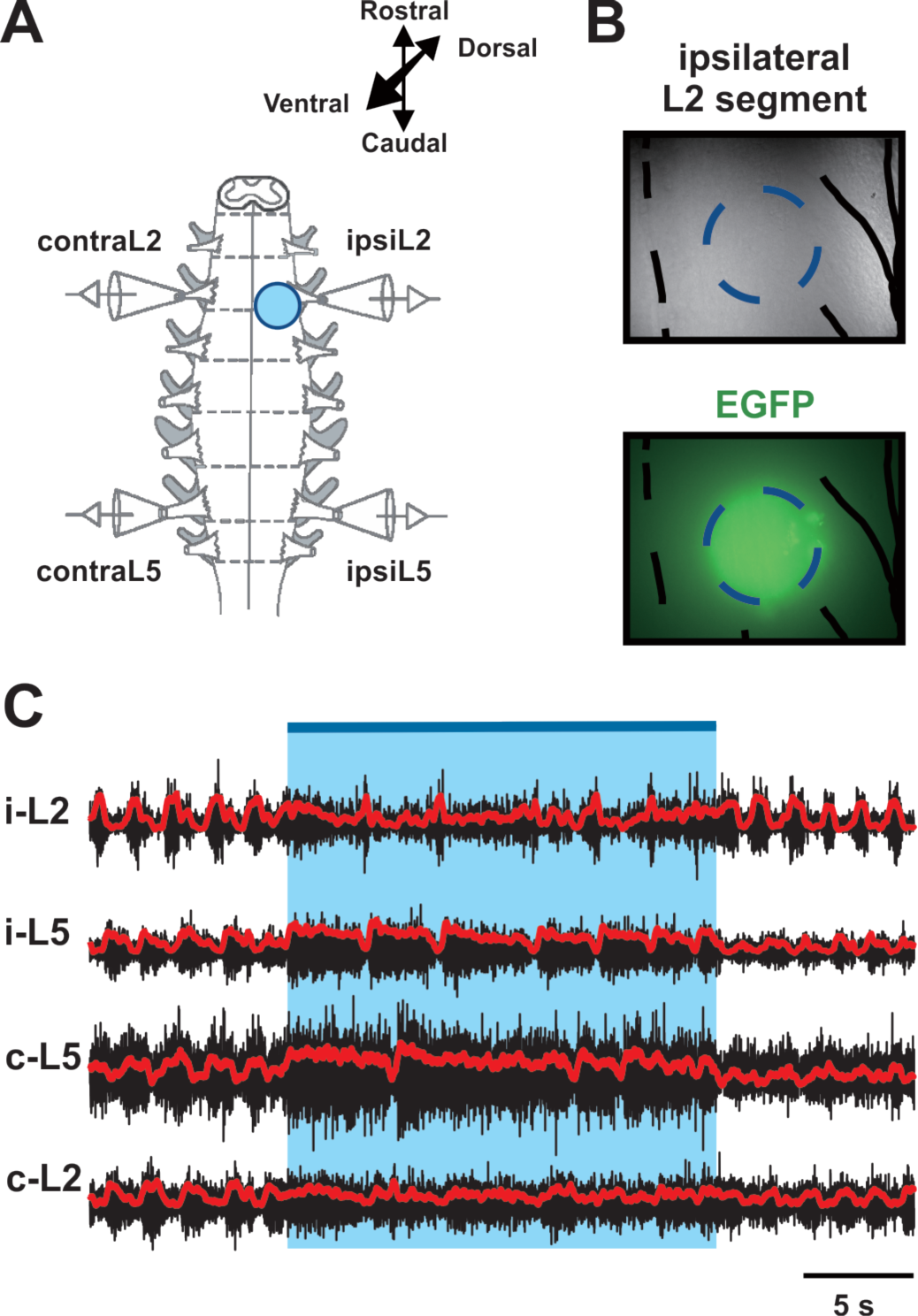
Biased optical activation of V3 interneurons in spinal segment L2. (A) Illustration of experimental setup for recording motor activity evoked by optical stimulation on the left spinal segment L2 from ventral side during drug-evoked fictive locomotion. (B) Image of the optical stimulation area. Black dashed line represents the midline of spinal cord. Blue dashed circle illustrates the light illuminated region. Black line on the right shows the frame of Nerve L2 and lateral edge of the spinal cord. (C) Recording trace during optical stimulation. Biased optical activation of V3 interneurons in spinal segment L2 leads to asymmetrical left-right extensor motor activity. Blue bar and area indicate the stimulation period. Red lines are rectified and smoothed traces of the original ENG recordings (black).

Under these experimental conditions, we found that optical activation of V3 interneurons on one side of the cord significantly prolonged L5 burst durations and step cycles on both sides (Figures 4C and 5A2,B; P < 0.0001), while L2 burst durations were not significantly affected (Figure 5A1). However, the contralateral L5 burst durations were influenced more strongly than those of ipsilateral L5 (Figures 6). Consequently, the step-cycle period also changed more strongly on the contralateral side than the ipsilateral side (Figure 5B), which led to a left-right asymmetric activity with more bursts on the ipsilateral than on the contralateral side. Emergence of additional bursts in an integer relationship is a fundamental property of (weakly-)coupled oscillators, with asymmetric drive (Pikovsky et al., 2001; Pikovsky and Rosenblum, 2003; Rubin et al., 2011). Thus, these results suggest that tonically activated V3 neurons mainly affect (and slow down) the contralateral CPG.

**Figure 5.**
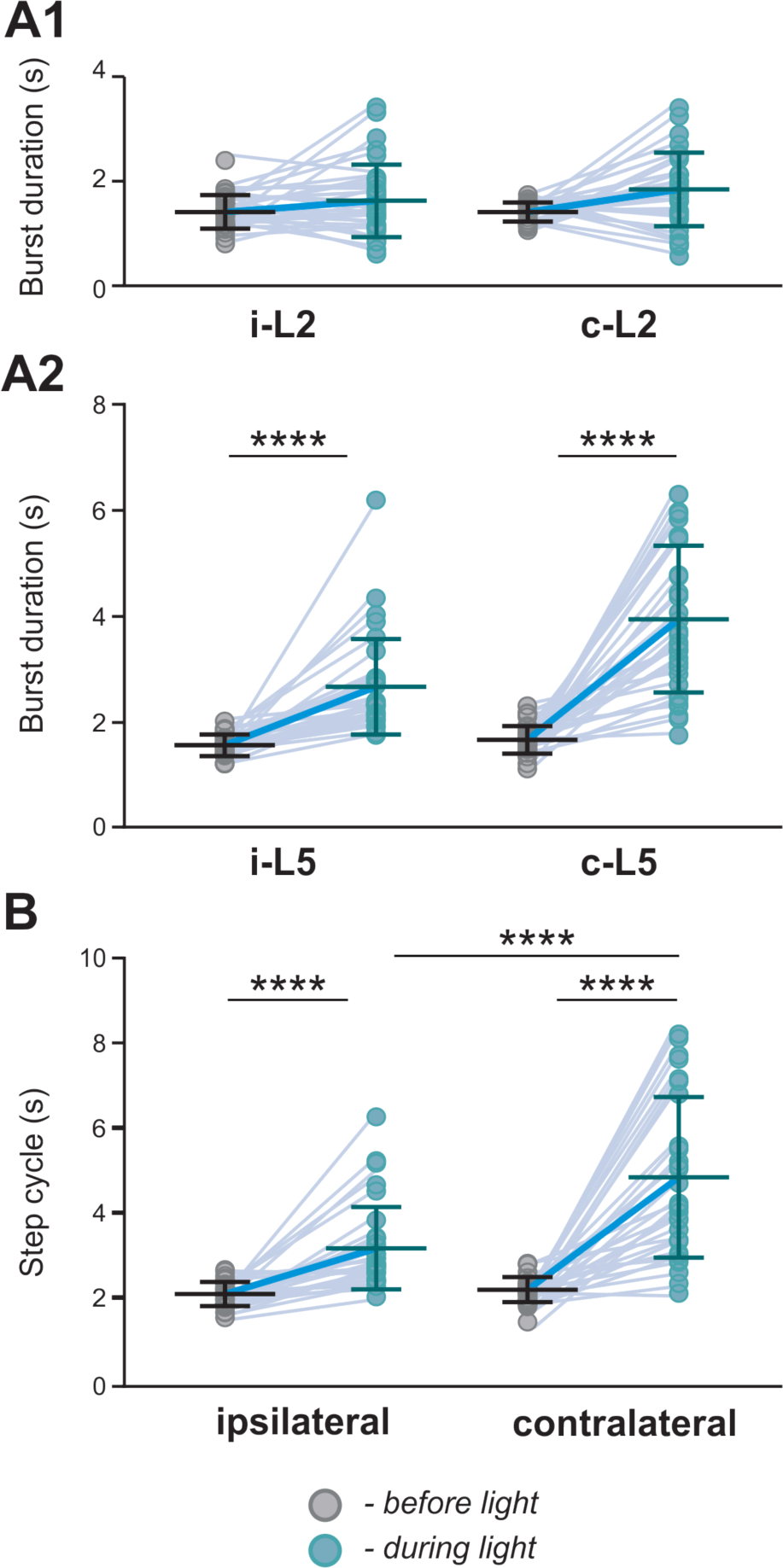
Changes in ENG characteristics by biased optical stimulation of V3 interneurons in spinal segment L2 during drug-induced locomotor-like activity. Average of burst duration of L2 (A1) and L5 (A2) during drug-induced locomotor-like activity (n = 33) before (grey dots) and during (blue dots) optical stimulation in ipsilateral and contralateral cord. (B) The average of step cycle period of ipsilateral and contralateral L5 activities during drug-induced locomotor-like activity (n = 33) before (grey dots) and during (blue dots) optical stimulation. Light blue lines connecting two dots represent change in one cord. Thick, dark blue line represent the average of change. **** indicates P <0.0001.

**Figure 6.**
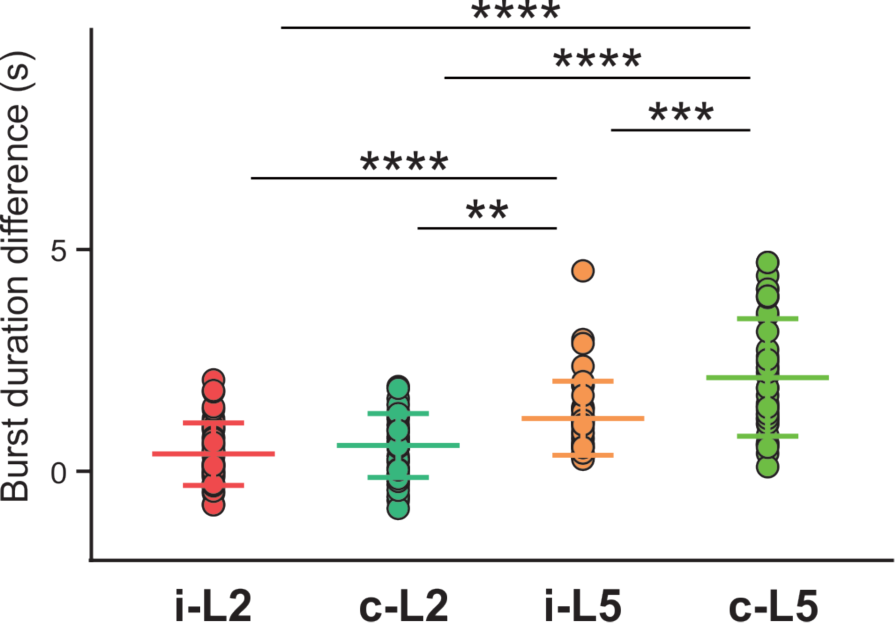
Average of locomotor burst duration difference between before and during biased optical stimulation. ** indicates 0.001 < P < 0.01, *** indicates 0.0001 < P <0.001, **** indicates P <0.0001; n = 33.

We also noticed a dose-response relationship between stimulation intensity and the prolongation of the contralateral cycle period and L5 burst duration (Figure 7A-C). At the highest applied light intensity, the rhythm of the contralateral cord can be suppressed, resulting in almost sustained extensor activity (Figure 7C). To further evaluate the relation between the asymmetric changes of the fictive locomotor activity in two sides of the spinal cord and the imbalanced activation intensity of V3 neurons, we systematically manipulated the focal size and light intensity, as described in the methods, to study the response to three levels of stimulation (low, medium and high intensity). Interestingly, when we plotted the changes of the burst duration and step cycle by the light activation against the stimulation intensity, we found that the L5 burst duration on the contralateral side to the light stimulation showed positive linear correlation to the optical stimulation intensity (Figure 8A1,A2; R^2^= 0.5081, P < 0.0001), but not the ipsilateral L5 (R^2^ = 0.007813) or both L2s (R^2^ = 0.003418, R^2^ = 0.1186 for ipsilateral and contralateral sides, respectively). In turn, the changes of step cycle of contralateral locomotor activities also showed a positive linear correlation with the optical stimulation intensity (Figure 8B; R^2^= 0.4842, P < 0.0001). This result indicates that the asymmetric response is dependent on the activation of V3 neurons on one side of the spinal cord.

**Figure 7.**
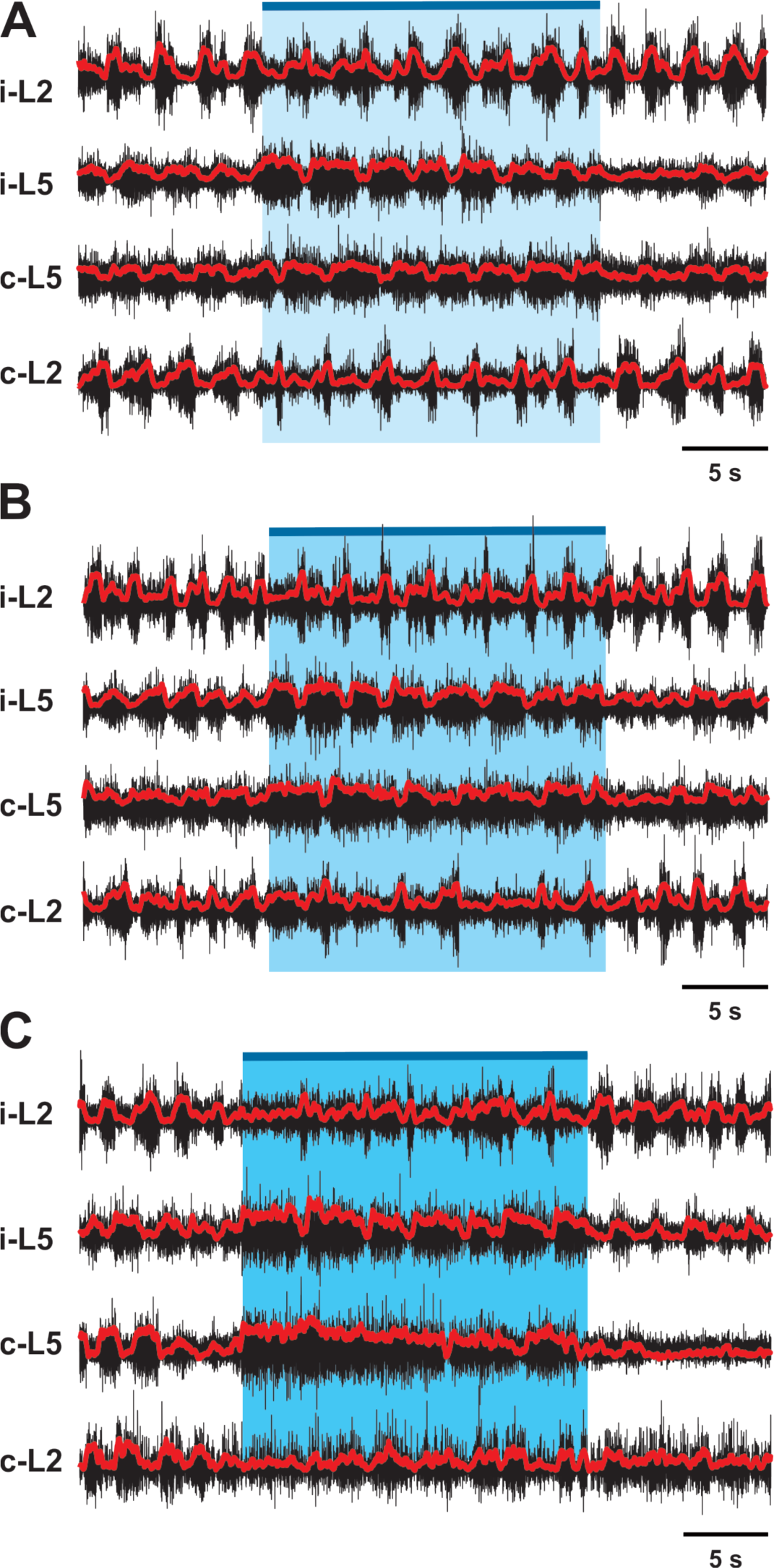
V3 activated contralateral extensor motor activity is correlated with optical stimulation intensity. Representative traces of locomotor activity induced by drug (5-HT 8μM, NMDA 8 μM) in *Sim1*^*Cre/*+^*; Ai32* cord at low (A), medium (B) and high (C) light intensity. Blue bar on the top of each trace and shaded blue area represent the optical stimulation period. The brightness of blue shaded area indicates light intensity.

**Figure 8.**
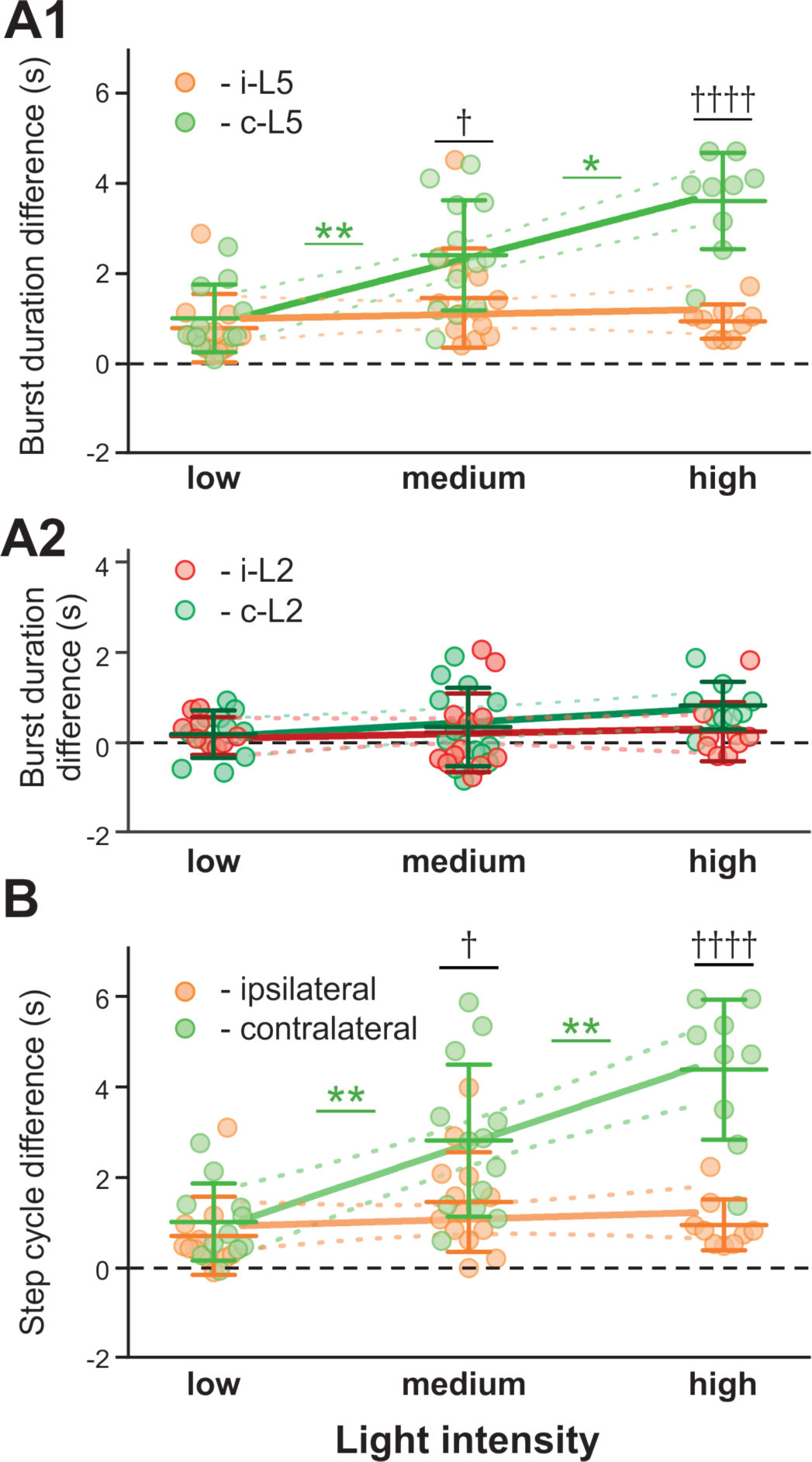
Dependence of ENG characteristics on light intensity during biased optical stimulation. Average of burst duration difference of L5 (A1) and L2 (A2) between before and during optical stimulation under low, medium and high stimulation intensity during drug-induced locomotor-like activity (low n = 11, medium n = 13, high n = 9). * indicates 0.01 < P <0.05, ** indicates 0.001 < P < 0.01. † indicates 0.01 < P <0.05, †††† indicates P <0.0001. Linear regression slope is orange for ipsilateral L5 and light green for contralateral L5. Slope of contralateral L5 burst duration and step cycle difference are significantly non-zero. (B) Step cycle period difference between ipsilateral and contralateral L5 under low, medium and high stimulation intensity during drug-induced locomotor-like activity (low n = 11, medium n = 13, high n = 9). †††† indicates P <0.0001, ** indicates 0.001 < P < 0.01.

Together, our experiments showed that the activation of V3 neurons increased extensor activity, and prolonged burst and step cycle duration more predominantly on the contralateral side with a smaller effect on the ipsilateral side. These resulted in left-right asymmetric rhythmic activity with lower-frequency bursting on the contralateral than on the ipsilateral side, suggesting that V3 neurons mainly affect the extensor activity of the contralateral CPG.

### 2.3 Modeling left-right interactions between rhythm generator

To further delineate the function and connectivity of the spinal V3 interneurons involved in left-right coordination, we developed a computational model of the lumbar locomotor circuitry. We built upon our previous model (Shevtsova et al., 2015) with the assumption that V3 neurons slow down the contralateral rhythm by exciting the extensor centers of the contralateral rhythm generators and transsynaptically inhibiting the flexor centers. Our goal was to update the model so that it could reproduce the effect of bilateral and unilateral stimulation of V3 neurons revealed in the above experiments without disrupting its ability to reproduce previous experimental findings concerning the effects of ablation of V0_V_, V0_D_ and all V0 CINs on the left-right coordination (Talpalar et al., 2013).

#### 2.3.1 Model schematic

The updated model consisted of two rhythm generating networks (RGs), one for each side of the spinal cord (Figure 9). Each RG included two excitatory populations, representing flexor (F) and an extensor (E) RG centers that mutually inhibited each other through populations of inhibitory interneurons (InF and InE). Similar to our previous model (Shevtsova et al., 2015), all neurons in both RG centers included a persistent (slowly-inactivating) sodium current, allowing them to intrinsically generate rhythmic bursting. Because of the mutual excitation between the neurons within each center, that synchronized their activity, each center could intrinsically generate a population rhythmic bursting activity. However, according to the setup of initial neuronal excitability, under normal conditions, the extensor centers, if uncoupled, expressed sustained activation and the rhythmic activity of each RG were defined by the activity of flexor centers, which then provided rhythmic inhibition of the corresponding extensor centers via the InF populations (see Figure 9 and Rybak et al., 2015; Shevtsova et al., 2015; Shevtsova and Rybak, 2016; Zhong et al., 2012). Interactions between F and E centers of the left and right RGs were mediated by populations of V3, V0_V_ and V0_D_ CINs (Figure 9). Drug-induced fictive locomotion was modeled by an unspecific increase of the excitability of all neurons in the network through a depolarization of the leakage reversal potentials in each neuron. The drug concentration was defined by the parameter α (see Material and Methods).

**Figure 9.**
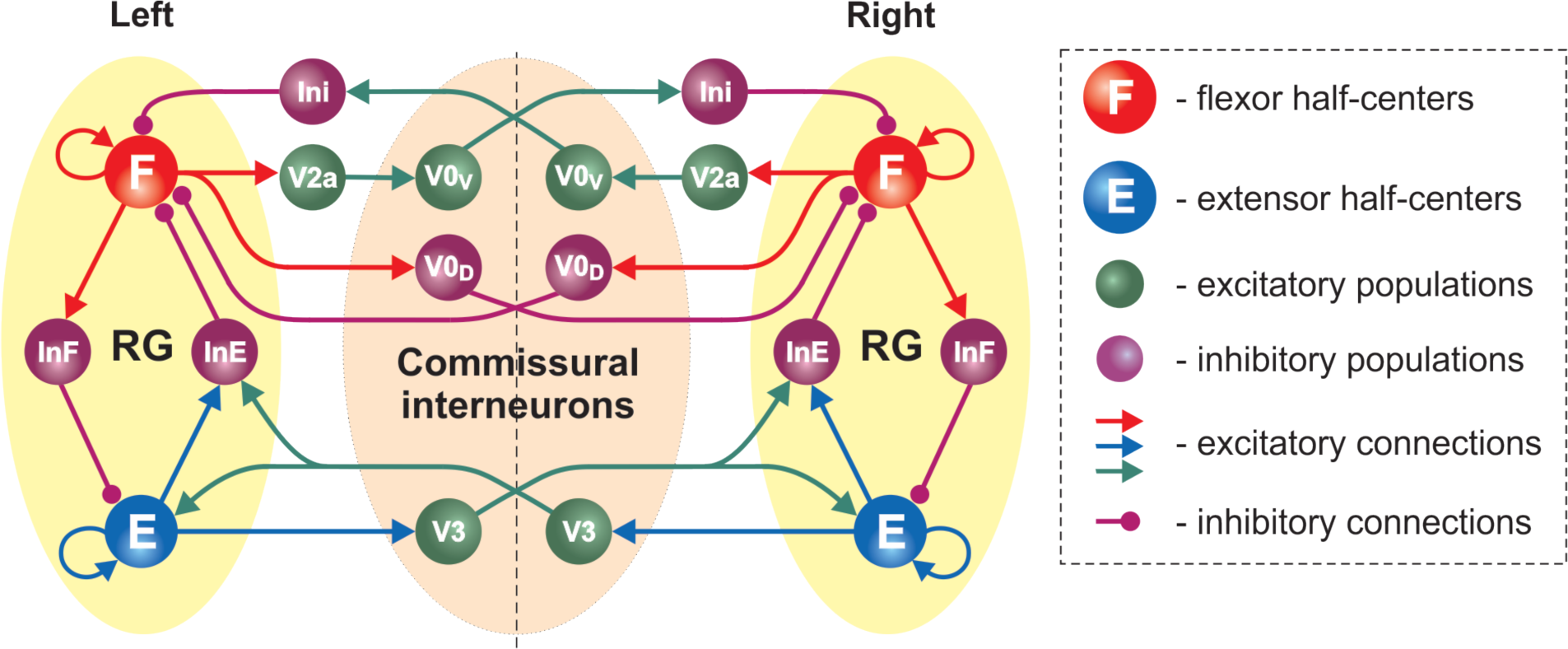
Model schematic of the bilateral spinal circuits consisting of two interconnected rhythm generators. Spheres represent neural populations and lines the synaptic connections between them. The rhythm generator (RG) in each side includes flexor and extensor centers (F and E, respectively) interacting via the inhibitory InF and InE populations. The left and right RGs interact via commissural interneurons (V0_D_, V0_V_, and V3).

In the present model, the organization of interactions between left and right RGs mediated by V0_V_ and V0_D_ populations of CINs followed that of our previous models (Ausborn et al., 2019; Danner et al., 2016, 2017; Shevtsova et al., 2015; Shevtsova and Rybak, 2016). Specifically (see Figure 9), the populations of inhibitory V0_D_ CINs provided direct mutual inhibition between the left and right flexor centers of the RGs, while the populations of excitatory V0_V_ CINs mediated mutual inhibition between the same flexor centers through oligosynaptic pathways (each V0_V_ population received excitation from the ipsilateral flexor center through a local population of V2a neurons and inhibited the flexor center of the contralateral RG through a population of local inhibitory neurons, Ini). Both V0_V_ and V0_D_ pathways ensured left-right alternation.

The organization of V3 CIN pathways in the present model differed from the previous models and was constructed to fit our experimental results (see section 2.1 and 2.2). Based on these results we suggested that V3 populations mediate mutual excitation between the extensor centers of the left and right RGs (Figure 9) and promoted inhibition of the contralateral flexor centers.

Each population of neurons in our model (Figure 9) consisted of 50–200 neurons (Table 1). All neurons were modelled in the Hodgkin-Huxley style (for details see Materials and Methods, section 4.4 and Table 1). Heterogeneity within the populations was ensured by randomizing the baseline value for leakage reversal potential and initial conditions for the values of membrane potential and channel kinetics variables. Connections between the populations were modelled as sparse random synaptic connections. Model equations and simulation procedures are listed in Material and Methods. Population specific parameters are listed in Table 1 and connection weights and probabilities are specified in Table 2.

**Table 1.** Number of neurons and neuron parameters in different populations.

| Neuron type | N, number of neurons | $\bar{g}_{Na}$, mS/cm<sup>2</sup> | $\bar{g}_{NaP}$, mS/cm<sup>2</sup> | $\bar{g}_K$, mS/cm<sup>2</sup> | $g_L$, mS/cm<sup>2</sup> | $\bar{E}_{LO}$, mV |
| --- | --- | --- | --- | --- | --- | --- |
| F | 200 | 25 | 0.75(±0.00375) | 2 | 0.07 | -67(±0.67) |
| E | 100 | 25 | 0.75(±0.00375) | 2 | 0.07 | -60(±0.6) |
| InF | 100 | 10 |  | 5 | 0.1 | -67(±1.34) |
| InE | 100 | 10 |  | 5 | 0.1 | -72(±1.44) |
| V0D | 50 | 10 |  | 5 | 0.1 | -68(±2.04) |
| V2a | 50 | 40 |  | 5 | 0.8 | -60.5(±1.21) |
| V0V | 50 | 10 |  | 5 | 0.1 | -62(±1.24) |
| Ini | 50 | 10 |  | 5 | 0.1 | -64(±1.28) |
| V3 | 100 | 10 |  | 5 | 0.1 | -68(±2.04) |

**Table 2.** Average weights 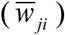 and probabilities (*p*) of synaptic connections.

| Source population | Target populations<br>($\bar{w}_{ji}$, probability of connection $p$) |
| --- | --- |
| i-F | i-F(0.009, $p=0.1$);<br>i-InF(0.01, $p=0.1$);<br>i-V2a(0.01, $p=0.1$);<br>i-V0D(0.06, $p=0.1$) |
| i-E | i-E(0.018, $p=0.1$);<br>i-InE(0.12, $p=0.1$);<br>i-V3(0.2, $p=0.05$) |
| i-InF | i-E(-0.15, $p=0.1$) |
| i-InE | i-F(-0.3, $p=0.1$) |
| i-Ini | i-F(-0.5, $p=0.1$) |
| i-V2a | i-V0v(1.5, $p=0.1$) |
| i-V0v | c-Ini(1.5, $p=0.1$) |
| i-V0D | c-F (-0.18, $p=0.1$) |
| i-V3 | c-E(0.05, $p=0.05$);<br>c-inE(1, $p=0.05$) |

Prefixes i- and c- indicate ipsi- and contralateral populations.

#### 2.3.2 The model exhibits characteristic features of drug-induced fictive locomotion and frequency-dependent changes of left-right coordination following removal of V0 commissural interneurons

First, we characterized the model performance under normal conditions by simulating drug-induced fictive locomotion (Figure 10A-C). The model exhibited alternation between the rhythmic activities of the flexor and extensor RG centers on each side as well as alternation between the activities of the left and right RGs. By increasing α (simulating an increase in the drug concentration) the burst frequency increased (Figure 10A-C). The frequency increase was accompanied by an asymmetric decrease of the burst durations: extensor burst durations decreased more than flexor burst durations. At all frequencies, left-right alternation was maintained. Thus, the model reproduced the main characteristics of drug-induced fictive locomotion in mice.

**Figure 10.**
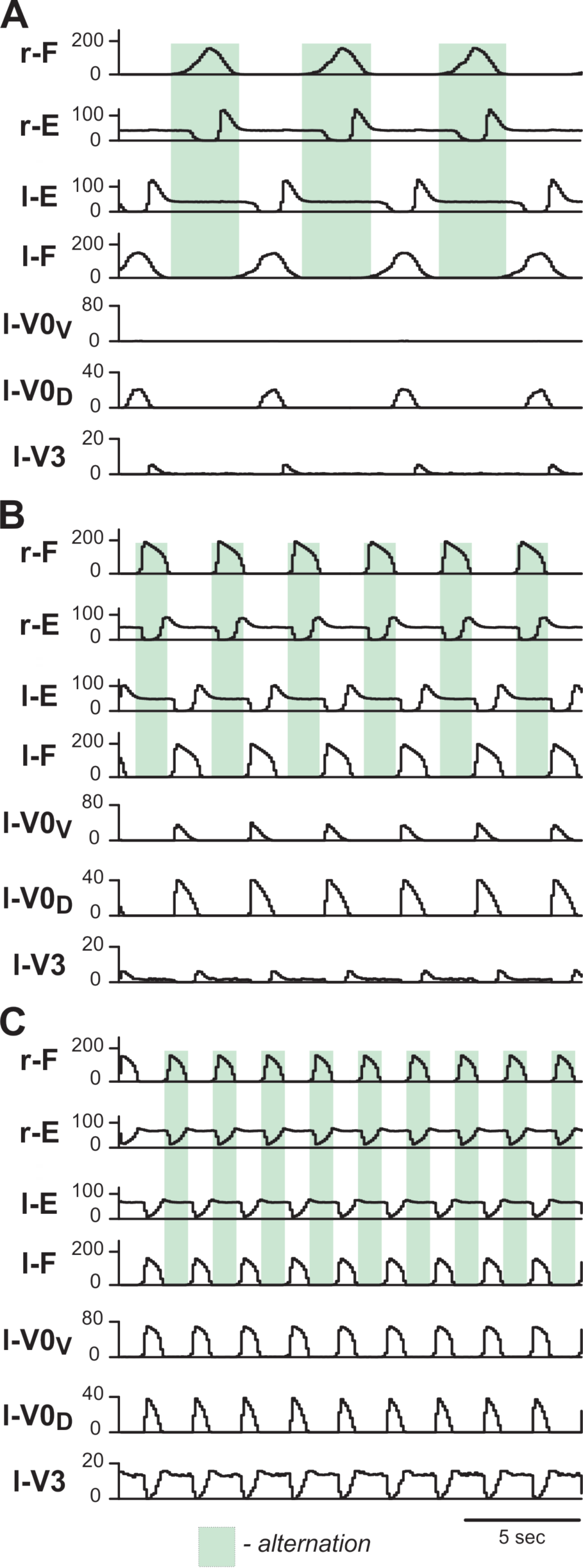
Model performance under normal conditions. Integrated population activities of flexor (F) and extensor (E) rhythm generator centers and left commissural interneurons are shown at low (A; α=0.01), medium (B; α=0.03) and high excitation levels (C; α=0.09). Activities of populations in this and following figures are shown as average histograms of neuron activity [spikes/(N × s), where N is a number of neurons in population; bin = 100 ms]. Shaded green areas show intervals of right flexor activities. r-: right; l-: left.

To test whether the model is still consistent with the frequency-dependent changes in left-right coordination following the removal of V0 CINs (Talpalar et al., 2013), we simulated the selective removal of V0_V_, V0_D_ or both V0 CIN populations by setting all connection weights from the selected types of neurons to 0. Removal of V0_V_ CIN populations did not change left-right alternation at low locomotor frequencies (Figure 11A1), but demonstrated left-right synchronized activity at high oscillation frequencies (Figure 11A2), Removal of V0_D_ CINs had the opposite effect: left-right synchronization occurred at low frequencies (Figure 11B1), while left-right alternation was maintained at high frequencies (Figure 11B2). Finally, removal of both types of V0 CIN populations led to left-right synchronization at all frequencies (Figure 11C1,C2). Thus, similar to the previous model (Shevtsova et al., 2015) the present model was able to reproduce the experimental results on the speed-dependent role of V0_V_ and V0_D_ in support of left-right alternation (Talpalar et al., 2013).

**Figure 11.**
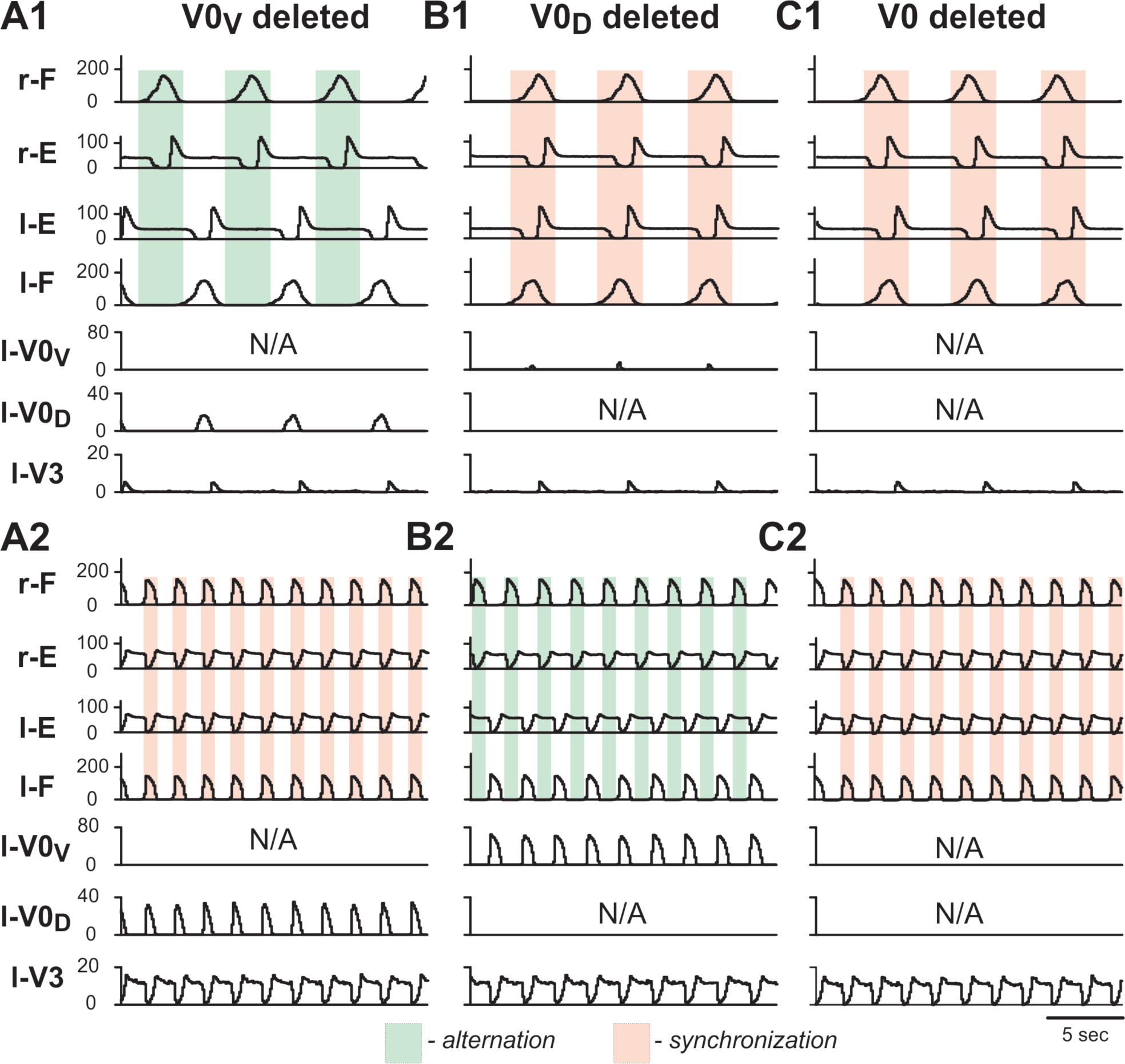
Model performance after removal of V0_V_, V0_D_ and all V0 commissural interneurons. Integrated population activities of flexor and extensor rhythm generator centers (F and E, respectively) and left commissural interneurons are shown after removal of V0_V_ (A1, A2), V0_D_ (B1, B2), and all V0 (C1, C2), at a low (A1, B1, C1; α=0.01) and a high (A2, B2, C2; α=0.08) excitation level and oscillation frequency. Shaded areas show intervals of right flexor activities. Green indicates left-right alternation and pink left-right synchronization. r-: right; l-: left.

The main difference between the previous models and the present one is in the role of V3 CINs and organization of their connections: in the previous models V3 CINs mediated mutual excitation between the flexor centers of the left and right RGs (Rybak et al., 2015; Shevtsova et al., 2015; Shevtsova and Rybak, 2016), whereas in the present model they mediate mutual excitation between the extensor centers (Figure 9). In both cases, however, these CINs promote left-right synchronization when mutual inhibition between the left and right RGs mediated by V0 CINs becomes weaker or has been partly or fully eliminated genetically.

#### 2.3.3 The model reproduces deceleration of the rhythm by tonic stimulation of V3 neurons

To simulate bilateral optogenetic stimulation of V3 CINs (section 2.1), we incorporated the channelrhodopsin ionic current, *I*_ChR_, in V3 neurons (see Material and Methods, section 4.4.3), which was activated in all V3 neurons for 15 seconds during ongoing locomotor activity. The results of these simulations with progressively increased stimulation intensity are shown in Figure 12. At any value, the applied stimulation increased the firing rate of active V3 neurons and recruited new neurons that were silent before stimulation (Figure 12A2-C2, A3-C3). Immediately with its onset, V3-stimulation increased the cycle period and reduced the burst frequency of both RGs (Figure 12A1,B1). The frequency reduction was mainly caused by a prolongation of the extensor burst duration. Flexor-extensor and left-right alternation were preserved during this stimulation. With increasing value of stimulation, the frequency was progressively decreased (Figure 12A1,B1). At high stimulation values, the rhythm could be stopped, resulting in sustained activation of both extensor centers and suppression of both flexor centers (Figure 12C1). Once the stimulation was stopped, the model exhibited a short transitional period (one or two cycles) and then returned to the same burst frequency and pattern as were expressed before stimulation. Altogether these results show that the model closely reproduces our experimental findings of optogenetic stimulation of V3 neurons during drug-induced fictive locomotion when stimulation was applied at the midline and equally affected left and right V3 neurons (section 2.1).

**Figure12.**
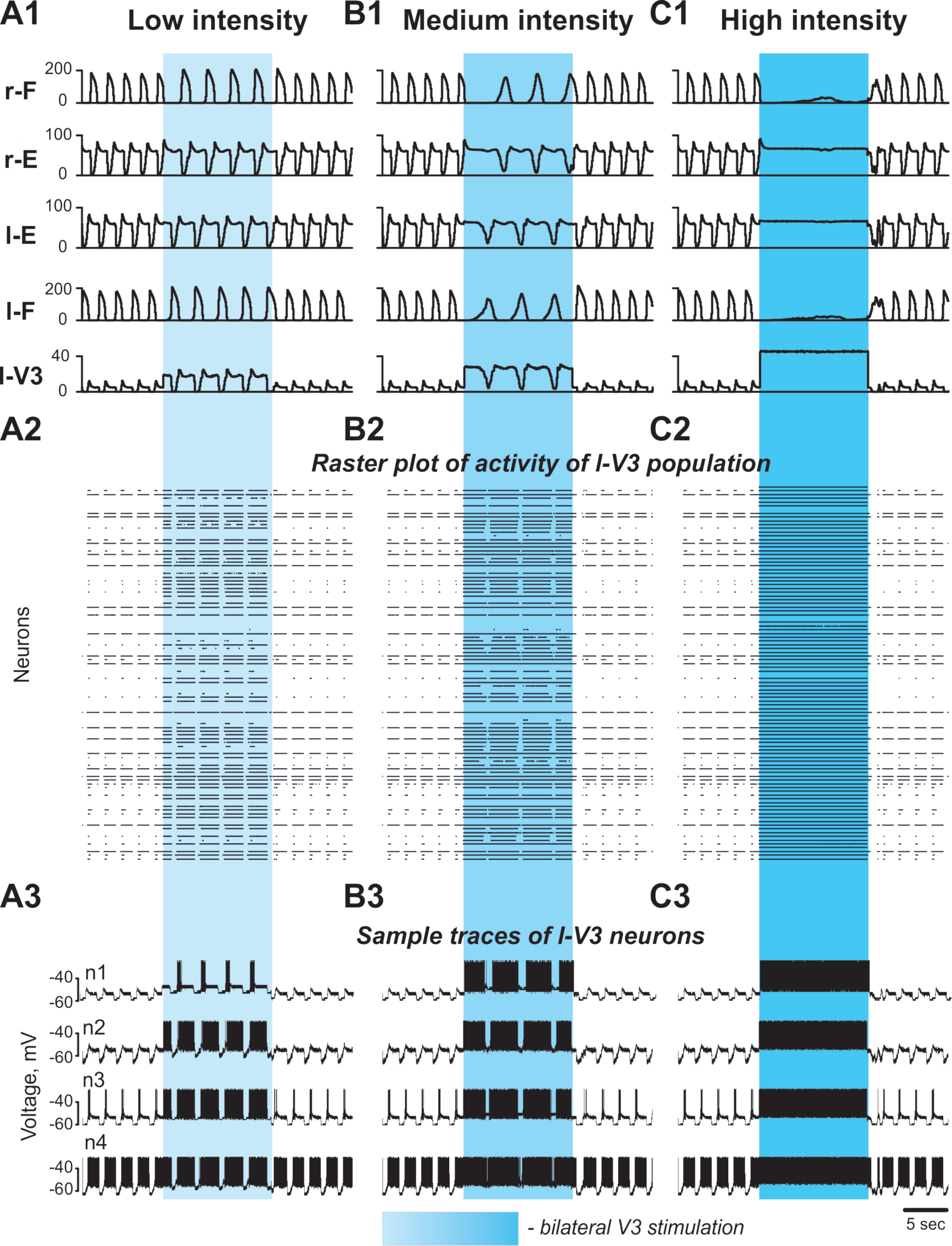
Effect of bilateral stimulation of V3 commissural interneurons during fictive locomotion in the model. (A1-A3) Low V3-stimulation intensity (*g*_chR_ =0.2). (B1-B3) Medium V3-stimulation intensity (*g*_chR_ =0.3). (C1-C3) High V3-stimulation intensity (*g*_chR_ =0.5). Stimulation was applied to all V3 neurons (on both sides of the cord). For all stimulations fictive locomotion was evoked at α=0.06. (A1, B1, C1) Integrated population activities of the flexor and extensor rhythm generator centers and the left V3 population. (A2, B2, C2) Raster plots of spikes elicited by neurons in the left V3 population. (A3, B3, C3) Traces of the membrane potential of sample neurons in the left V3 population with different excitability. Shaded blue areas show the interval when V3 stimulation was applied. r-: right; l-: left.

The reduction of the burst frequency when V3 neurons were stimulated occurred because both (left and right) extensor centers were activated by V3 neurons and they both provided an additional inhibition to the corresponding flexor centers through the corresponding inhibitory populations (InE), which reduced the average excitation of the flexor centers (Figure 9). In addition, activated V3 neurons provided direct excitation of the contralateral InE populations inhibiting the corresponding flexor centers. Note that the frequency of persistent sodium current-dependent oscillations positively correlates with the average excitation of a population of neurons with this current and mutually excitatory interconnection (Butera et al., 1999; Rybak et al., 2004, 2015). Since the rhythm in our model was generated by flexor centers, the reduction of their excitation during V3 neuron activation led to the reduction of oscillation frequency generated in both RGs. Furthermore, the reciprocal excitation of the extensor centers through V3 CINs created a positive feedback loop that amplified the firing rates of its constituent neurons and consequently the net inhibition exerted on the flexor centers.

#### 2.3.4 The model reproduces asymmetric changes of the locomotor rhythm by unilateral stimulation of V3 neurons

To simulate the effects of unilateral activation of V3 neurons during locomotor activity (see section 2.2), we activated *I*_ChR_ in all V3 neurons located on one side of the cord. At a low value of unilateral activation of ipsilateral V3 population (Figure 13A), the extensor burst durations and the cycle periods increased on both sides, while the flexor-extensor and left-right alternation remained unchanged, which was similar to the effect of bilateral stimulation. With increasing value of stimulation (Figure 13B,C), the rhythm of the contralateral circuits was progressively slowed down, and additional bursts appeared ipsilateral to the stimulation. Figure 13B shows a stable 2-to-1 relationship between the number of burst ipsilateral and contralateral to the stimulation. In Figure 13C one can see both 2-to-1 and 3-to-1 relationships. In all cases, reciprocity between the flexor centers maintained. Our simulations shown in Figure 13A,B qualitatively reproduced the experimental results concerning the response of the fictive locomotor pattern to the weak and medium unilateral optical stimulation of V3 neurons on (Figure 7A,B) where one can also see unequal number of flexor bursts on ipsi- and contralateral sides.

**Figure 13.**
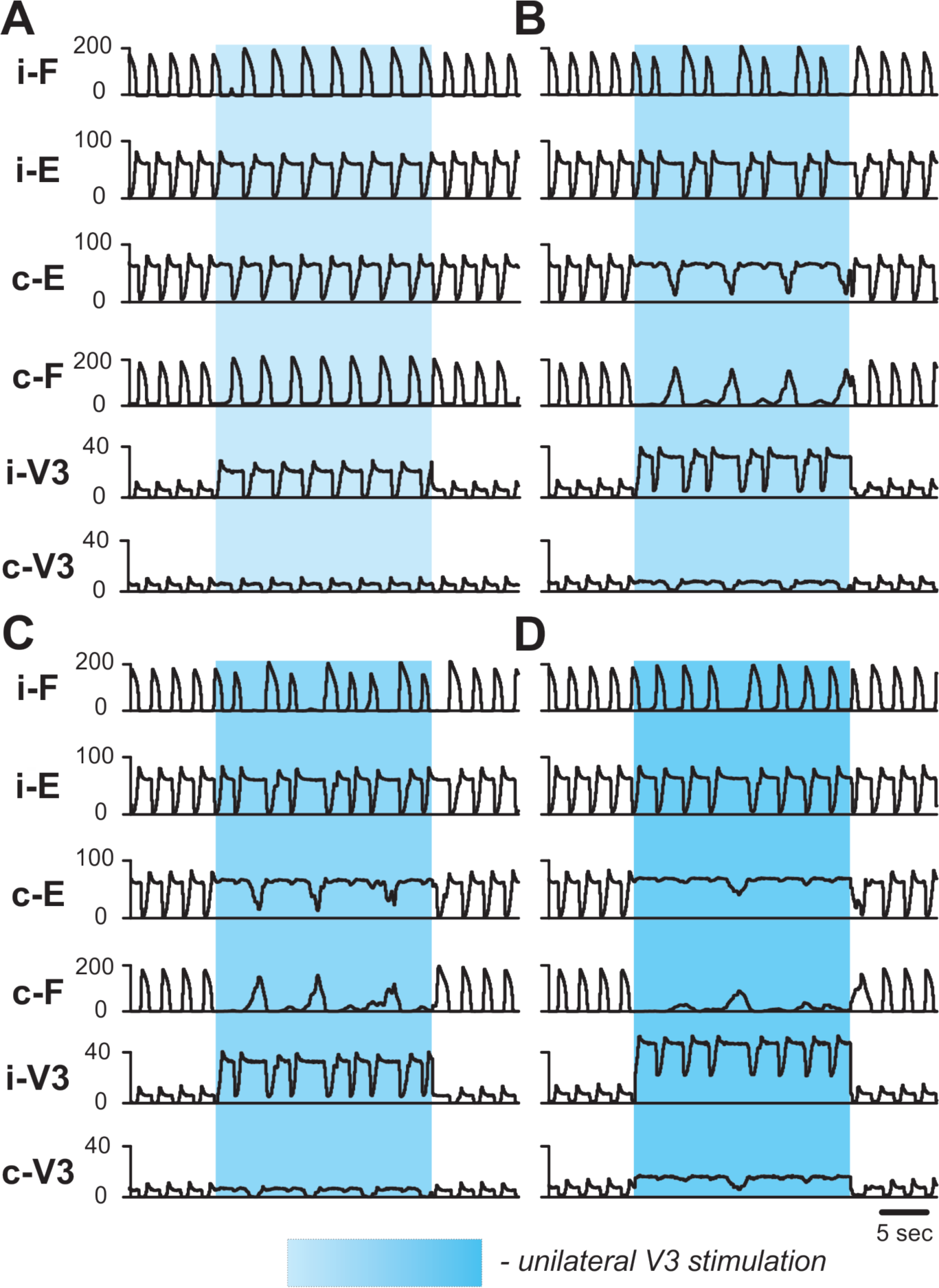
Effect of unilateral stimulation of V3 commissural interneurons during fictive locomotion in the model. Fictive locomotion was evoked at α=0.065. All panels show integrated population activities of the flexor (F) and extensor (E) rhythm generator centers and the V3 populations ipsilateral (i-) and contralateral (c-) to the stimulation. Shaded blue areas show the interval when V3-stimulation was applied. Four different stimulation intensities are shown: (A) *g*_chR_ = 0.22; (B) *g*_chR_ = 0.33; (C) *g*_chR_ = 0.35; (D) *g*_chR_ = 0.5 in ipsilateral and *g*_chR_ = 0.1 in contralateral V3 population. i-: ipsilateral; c-: contralateral.

To simulate the effects of high intensity and larger focal area unilateral stimulation, an additional weak stimulation of the contralateral V3 neuron population was applied (Figure 13D). In this case, the rhythmic activity on the ipsilateral side slowed down during the stimulation and the contralateral extensor centers became constantly active, while irregular low-amplitude activity of the contralateral flexor centers occurred. This behavior of the model was similar to the results of experimental studies at a high intensity unilateral stimulation when the rhythm of the contralateral cord was suppressed, resulting in almost sustained extensor activity (Figure 7C). Based on our simulation results, we suggest that at high stimulation intensities the stimulation partly affected V3 neurons on the side contralateral to stimulation.

## 3 Discussion

In the current study, we combined experimental studies with computational modeling to investigate the potential roles that V3 interneurons play in the spinal locomotor network. Using optogenetic approaches, we were able to selectively activate V3 interneurons at different regions of the spinal cord during drug-evoked fictive locomotion, and our computational model of locomotor circuits was able to qualitatively reproduce the experimental results. Our study revealed that lumbar V3 interneurons strongly enhance the contralateral extensor activity and regulate the frequency of the locomotor oscillations. We suggest that spinal V3 CINs mediate mutual excitation between the extensor centers of the left and right rhythm generators in the lumbar spinal cord, which might support the synchronization of left-right activity under certain conditions during locomotion. In addition, we show that unilateral activation of V3 CINs can produce left-right asymmetrical rhythmic outputs, which provides a potential mechanism for the separate regulation of left-right limb movements, which may operate in certain conditions such as changing the direction of walking or during split-belt walking.

### 3.1 Optogenetic activation of V3 interneurons

Mapping the functional connectivity among neurons in the central nervous system is vital to understand the circuit logics underlying behavior. This is especially true within the spinal cord, which generates the rhythmic and patterned motor outputs necessary for coordinated movement (Danner et al., 2015; Grillner, 2006; Kiehn, 2006, 2011). However, uncovering the functional circuitry within the spinal cord was difficult due to a lack of clear anatomical organization compared to other brain regions. In the current study, novel optogenetic tools have enabled us to specifically (Dougherty et al., 2013; Hägglund et al., 2013; Levine et al., 2014), reversibly, and acutely regulate the activity of a molecularly-identified group of spinal neurons. By using *Sim1Cre/*+*; Ai32* mice (Chopek et al., 2018), we were able to target V3 interneurons on both or just one side of the isolated spinal cords allowing us to start dissecting the function of V3 CINs in spinal locomotor circuits.

Using whole-cell patch-clamp recordings, we could verify that all EYFP/ChR2-expressing cells in the lumbar spinal slices could be exclusively depolarized by blue fluorescent light and elicit spikes during periods of the light pulses (10–20 s). During our functional study, however, we had to keep the whole lumbar cord intact and deliver the LED light on the ventral surface of the isolated spinal cord, in which way, we would expect to activate V3 neurons that had cell bodies and terminals located or neurites passing the illuminated region that is relatively close to the ventral surface. Although we were not able to specify the particular group of V3s, which we might be activating, we kept the light intensity, size and position consistent to limit variability across experiments.

### 3.2 The V3 CINs provide excitation to the contralateral CPG extensor center

The identification of genetically defined interneuron populations has enabled us to characterize their specific functions in locomotor behaviors (Deska-Gauthier and Zhang, 2019; Gosgnach et al., 2017; Goulding, 2009). Until now, however, most experimental data were from genetically eliminating certain group of spinal interneurons without clear understanding of connectivity among the spinal interneurons in spinal circuits (Bellardita and Kiehn, 2015; Crone et al., 2008; Gosgnach et al., 2006; Lanuza et al., 2004; Talpalar et al., 2013; Zhang et al., 2008). In the case of V3 interneurons, genetic deletion of entire V3 population reduced the general robustness of the rhythmicity of the *in vivo* and *in vitro* locomotor activity and led to unstable gaits (Zhang et al., 2008). Our previous anatomical studies were only able to show that V3 cells had broad projections to contralateral motor neurons and other ventral spinal interneurons along the lumbar spinal cords, but we still do not know their precise anatomical position and their role in the CPG network (Blacklaws et al., 2015; Zhang et al., 2008). The current spatially controlled activation of V3 neurons was the first step to overcome this limitation to decipher their functional role in the spinal cord.

We found that the bilateral photoactivation of V3 neurons in the isolated spinal cord during fictive locomotion increased the intensity and duration of L5 bursts and prolonged the step cycles in all recorded lumbar roots whereas the duration of bursts in L2 roots was not significantly affected. These results suggested that, under our experimental condition, the photoactivated V3 interneurons predominantly excited the circuits responsible for extensor activities, which in turn reduced the frequency of fictive locomotion oscillations. This conclusion became more evident when we activated V3 interneurons only on one-side of the cord with varied light intensities. With such stimulations, we found that only the increase of burst duration of contralateral L5 were positively correlated to the intensity of the illuminating light, and at high intensity of unilateral photostimulation the contralateral L5 activity could express a sustained activity. More interestingly, such uneven regulation of contralateral extensor excitability by V3 CINs significantly decreased the bursting frequency of the contralateral ventral root output, which led to asymmetrical oscillation of the motor outputs on the two sides of the spinal cord. Thus, our experimental results support the conclusion that the lumbar V3 interneurons, or at least a sub-group of them, strongly regulate the extensor activity and step cycle duration on the contralateral side of the cord.

Similar to other genetically defined types of spinal interneuron populations (Bikoff et al., 2016; Deska-Gauthier and Zhang, 2019; Gosgnach et al., 2017), the V3 population is heterogeneous, consisting of sub-populations with different anatomical, physiological and molecular profiles (Borowska et al., 2013, 2015). In the present study, however, we did not consider different subtype of V3 neurons. When we delivered the LED fluorescent light on the ventral surface of the spinal cord, we most likely activated a mixed group of different V3 neuron types in the illuminated region. Nonetheless, this discovery has provided direct evidence that some subtypes of V3 neurons can regulate the rhythm and pattern of the locomotor output through manipulating the contralateral extensor circuits.

### 3.3 Coordination of left-right activities by V3 interneurons during locomotion

Until now, several types of CINs, including V0, V3, and dI6, have been identified in the ventral spinal cord (Andersson et al., 2012; Bellardita and Kiehn, 2015; Haque et al., 2018; Lanuza et al., 2004; Talpalar et al., 2013; Zhang et al., 2008). When each of these CIN types, except V3, was genetically deleted, the left-right hind-limb alternation was affected. Particularly, V0_V_ and V0_D_ CINs were shown to secure the speed-dependent left-right alternation at walking and trotting gaits without effects on hind-limb synchronization observed during gallop and bound (Bellardita and Kiehn, 2015; Talpalar et al., 2013). On the other hand, deletion of V3 CINs did not affect the left-right alternation of the limb movement during walk and trot (Zhang et al., 2008). Therefore, it has been suggested that V3 populations represent the CIN populations that mediate left-right synchronization in certain conditions (e.g. when some or all V0 CINs are genetically removed) and during gallop and bound. In our previous computational models (Danner et al., 2016, 2017; Rybak et al., 2013; Shevtsova et al., 2015; Shevtsova and Rybak, 2016), to perform this function the V3 CINs were assigned to explicitly mediate mutual excitation between the flexor centers of the left and right rhythm generators. However, our current experimental outcome contradicts this assumption, suggesting functional connection of V3 CINs to the contralateral extensor centers instead of the flexor centers. To match our experimental findings, we changed the connection of simulated V3 populations from the activation of contralateral flexor centers to the activation of contralateral extensor centers. With this change in the V3 connectivity we were able to qualitatively reproduce all our current and previous experimental results, including speed-dependent left-right alternation mediated by V0_V_ and V0_D_. This unification of the experimental and computational data confirmed that the proposed organization of our current computational model of spinal circuits is plausible, and that V3 interneurons can mediate the synchronization of left-right hind-limb activities through mutual excitation of extensor centers of the left and right spinal rhythm generators.

In addition, our experimental and modeling results suggest that V3 interneurons might play important functional roles in left-right coordination under some special conditions, such as changing the direction of movement (left or right turns, rotations) or on split-belt treadmills (Courtine and Schieppati, 2003a, 2003b; Forssberg et al., 1980; Hurteau and Frigon, 2018). In these situations, different speeds for left-right limbs are expected for some time intervals while maintaining stable locomotion. The precise mechanisms underlying such movements are still unknown, but it has been clearly indicated that the independent but closely coordinated CPG circuits in both sides of the spinal cord are involved. In our current experimental and modeling studies, we could produce separate locomotor outputs with two different oscillatory frequencies from the ventral roots on each side of the spinal cord by unilaterally activating V3 neurons. This result strongly suggested that V3 neurons could mediate the coordination of the left-right activity during asymmetrical movements, such as the split-belt walking, even though we still do not know whether and how this pathway is activated *in vivo*.

## 4 Material and Methods

### 4.1 Animals

The generation and genotyping of *Sim1*^*Cre/*+^ mice were described previously by Zhang et al. (2008). Ai32 mice were from the Jackson Laboratory (Stock No. 012569). It contains Rosa26-cytomegalovirus early enhancer element/chicken beta-actin promoter (CAG)-loxP–Stop codons–3x SV40 polyA–loxP (LSL)-channelrhodopsin2 (ChR2)-enhanced yellow fluorescent protein (EYFP)-woodchick hepatitis virus posttranscriptional enhancer (WPRE; RC-ChR2). *Sim1Cre/*+*; Ai32* mice generated by breeding these two strains expressed ChR2/EYFP fusion-protein in Sim1 expressing cells. All procedures were performed in accordance with the Canadian Council on Animal Care and approved by the University Committee on Laboratory Animals at Dalhousie University.

### 4.2 Electrophysiology

#### 4.2.1 Preparation

All experiments were performed using spinal cords from *Sim1Cre/*+*; Ai32* mice at postnatal day (P) P2-P3. The mice were anesthetized, and the spinal cords caudal to thoracic (T) 8 segments were dissected in an ice-cold oxygenated Ringer’s solution (111 mm NaCl, 3.08 mm KCl, 11 mm glucose, 25 mm NaHCO3, 1.25 mm MgSO4, 2.52 mm CaCl2, and 1.18 mm KH2PO4, pH 7.4). The spinal cord was then transferred to the recording chamber to recover at room temperature for 1 hour before recording in Ringer’s solution.

#### 4.2.2 Isolated whole-cord recordings

Electroneurogram (ENG) recordings of the (lumbar) L2 and L5 ventral roots were conducted using differential AC amplifier (A-M system, model 1700) with the band-pass filter between 300 Hz and 1 kHz. Analog signals were transferred and recorded through the Digidata 1400A board (Molecular Devices) under the control of pCLAMP10.3 (Molecular Devices). Fictive locomotor activity was induced by applying 5-hydroxytryptamine (5-HT, 8-10 μM) and NMDA (7-8 μM) in the Ringer’s solution.

#### 4.2.3 Optical stimulation of V3 interneurons

To activate ChR2 in V3 interneurons, 488 nm fluorescent light was delivered by Colibri.2 illumination system (Zeiss) through 10x or 20x 1.0 numerical aperture (NA) objectives mounted on an up-right microscope (Examiner, Zeiss) onto the ventral surface of the isolated spinal cord. The amount of light to activate ChR2 positive cells was adjusted by changing illuminated region size through the field diaphragm and/or the light brightness delivered by the Colibri (Zeiss, Figure 4A). We then manually adjust the field diaphragm and LED light intensity to tune the illuminated region between approximately one-third to a half of the spinal cord (Figure 4B) to regulate the number of V3 neurons being activated. For each experiment, we set the stimuli with largest focal area and 100% light intensity as the high intensity group, then reduced focal area as medium group, and then the smallest focal area and/or reduced light intensity as the low intensity group.

#### 4.2.4 Whole-cell patch-clamp recordings

The experimental procedures were detailed in (Borowska et al., 2013). Briefly, 300 uM slices from the spinal cord lumbar region (T13–L3) from P2-3 *Sim1Cre/*+*; Ai32* were prepared in an ice-cold oxygenated sucrose solution (3.5 mm KCL, 25 mm NaHCO3, 1.2 mm KH2PO4, 1.3 mm MgSO4, 1.2 mm CaCl2, 10 mm glucose, 212.5 mm sucrose, and 2 mm MgCl2, pH 7.4) on a vibratome (Vibratome 300, Vibratome). Slices were incubated in an oxygenated Ringer’s solution (111 mm NaCl, 3.08 mm KCl, 11 mm glucose, 25 mm NaHCO3, 1.25 mm MgSO4, 2.52 mm CaCl2, and 1.18 mm KH2PO4, pH 7.4) at room temperature for >30 min for recovery before recording. GFP fluorescence-positive and negative cells were visually identified using a 40× water-immersion objective (numerical aperture, 0.8) with the aid of a DAGE-MTI IR-1000 CCD camera.

Conventional whole-cell patch-clamp recordings were made in voltage- and current-clamp modes using a MultiClamp 700B amplifier (Molecular Devices). Analog signals were filtered at 10 kHz with the Digidata 1400A board (Molecular Devices) under control of pCLAMP10.3 (Molecular Devices). Patch-clamp recording pipettes with a resistance of 5–8 MΩ were filled with solution containing 128 mm K-gluconate, 4 mm NaCl, 0.0001 mm CaCl2, 10 mm HEPES, 1 mm glucose, 5 mm Mg-ATP, and 0.3 mm GTP-Li, pH 7.2 and the fluorescent light was delivered through the 40x objective.

### 4.3 Data analysis

All recorded traces were transferred to Spike2 (Version 7.09a, Cambridge Electronic Design). The data were analyzed using rectified and smoothed signals. The step cycle duration was calculated as the distance between the onset time of one burst and the onset of the next burst. The average of step cycle durations and burst durations of five consecutive cycles before the light-on and all the cycles during the light were used as a pair of data set of before and during light, respectively. The steps interrupted at the beginning or the end of light were excluded from the calculation.

Statistical analysis was performed in Prism7 (GraphPad Software, Inc.). Student’s paired t-test were used to compare the difference between the activity before and after the light. Linear contrasts were used to determine the relationship of the burst duration difference of activities in four nerves and step cycle difference in ipsilateral and contralateral side among three intensities. One-way ANOVA was used to determine the statistical significance among the burst duration of activity in four nerves. An α- error of P < 0.5 was regarded as significant. Data in the Results represent the mean ± SD.

### 4.4 Computational modelling

#### 4.4.1 Neuron model

All neurons were simulated in the Hodgkin-Huxley style as single-compartment models. The membrane potential, *V*, in neurons of the left and right flexor and extensor centers was described by the following differential equation

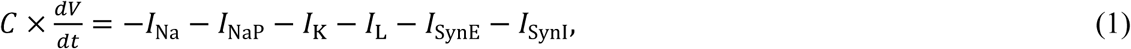

where *C* is the membrane capacitance and *t* is time.

In all other populations, the neuronal membrane potential was described as follows:

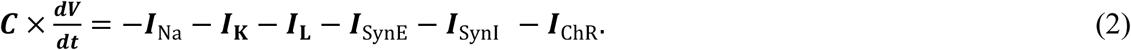

The ionic currents in Eqs. (1) and (2) were described as follows:

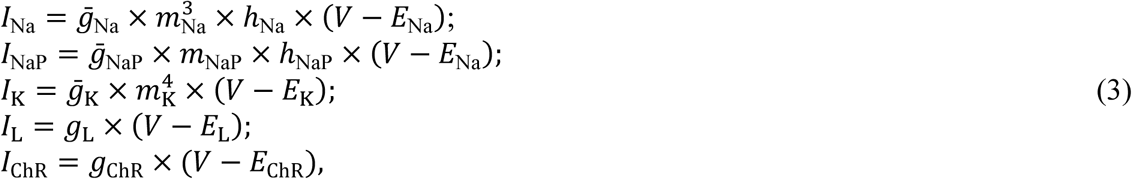

where *I*_Na_ is the fast Na^+^ current with maximal conductance 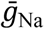; *I*_NaP_ is the persistent (slowly inactivating) Na^+^ current with maximal conductance 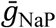 (present only in RG neurons); *I*_K_ is the delayed-rectifier K^+^ current with maximal conductance 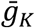; *I*_L_ is the leakage current with constant conductance *g*_L_; *I*_ChR_ is the channelrhodopsin current with the conductance *g*_ChR_ (present only in V3 neurons). *E*_Na_, *E*_K_, *E*_L_, and *E*_ChR_ are the reversal potentials for Na^+^, K^+^, leakage, and channelrhodopsin currents, respectively; variables *m* and *h* with indexes indicating ionic currents are the activation and inactivation variables of the corresponding ionic channels.

Activation *m* and inactivation *h* of voltage-dependent ionic channels (e.g., Na, NaP, and K) in Eq. (3) were described by the following differential equations:

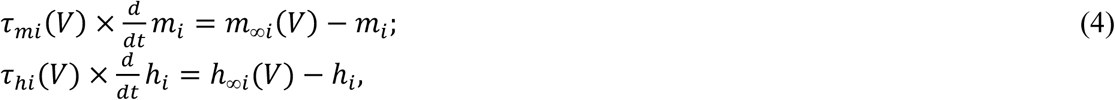

where *m*_∞*i*_(*V*) and *h*_∞*i*_(*V*) define the voltage-dependent steady-state activation and inactivation of the channel *i*, respectively, and *τ*_*mi*_(*V*) and *τ*_*hi*_(*V*) define the corresponding time constants. Activation of the sodium channels is considered to be instantaneous. The expressions for channel kinetics in Eq. (4) are described as follows:

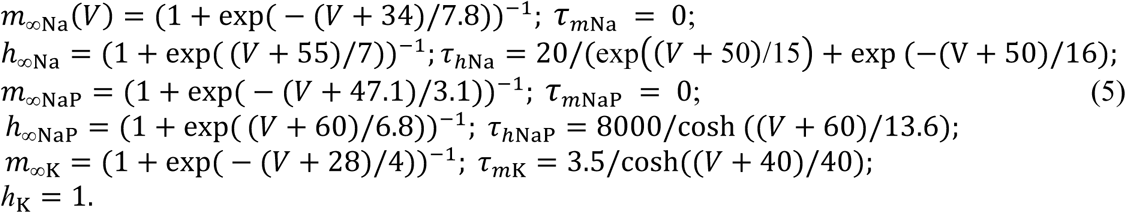

The maximal conductances for ionic currents and the leak reversal potentials, *E*_L_, for different populations are given in Table 1.

The synaptic excitatory (*I*_SynE_ with conductance *g*_SynE_ and reversal potential *E*_SynE_) and inhibitory (*I*_SynI_ with conductance *g*_SynI_ and reversal potential *E*_SynI_) currents were described as follows:

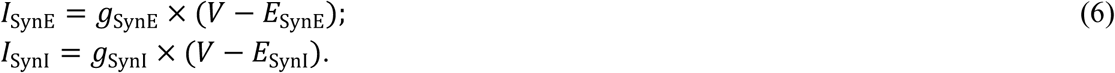

where *g*_SynE_ and *g*_SynI_ are equal to zero at rest and are activated by the excitatory or inhibitory inputs, respectively:

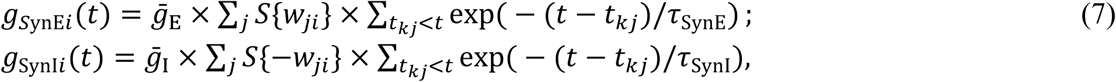

where *S*{*x*} = *x*, if *x* ≥ 0, and 0 if *x* < 0. Each spike arriving to neuron *i* in a target population from neuron *j* in a source population at time t_*kj*_ increases the excitatory synaptic conductance by 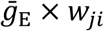 if the synaptic weight *w*_*ji*_> 0, or increases the inhibitory synaptic conductance by 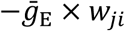 if the synaptic weight 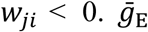 and 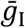 define an increase in the excitatory or inhibitory synaptic conductance, respectively, produced by one arriving spike at |*w*_*ji*_| = 1. *τ*_synE_ and *τ*_synI_ are the decay time constants for *g*_SynE_ and *g*_SynI_, respectively.

The following general neuronal parameters were assigned: *C* =1 μF·cm^−2^; *E*_Na_ = 55 mV; *E*_K_ = − 80 mV; *E*_ChR_ = -10 mV; *E*_SynE_ = −10 mV; *E*_SynI_ = −70 mV; 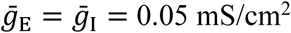;*τ*_synE_ = *τ*_synI_ = 5 ms.

#### 4.4.2 Neuron populations

Each neuron population in the model contained 50-200 neurons. The numbers of neurons in each population are shown in Table 1.

Random synaptic connections between the neurons of interacting populations were assigned prior to each simulation based on probability of connection, *p*, so that, if a population *A* was assigned to receive an excitatory (or inhibitory) input from a population *B*, then each neuron in population *A* would get the corresponding synaptic input from each neuron in population *B* with the probability *p*{*A, B*}. If *p*{*A, B*}<1, a random number generator was used to define the existence of each synaptic connection; otherwise (if *p*{*A, B*}= 1) each neuron in population *A* received synaptic input from each neuron of population *B*. Values of synaptic weights (*w*_*ji*_) were set using random generator and were based on average values of these weights 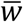 and variances, which were define as 5% of 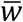 for excitatory connections 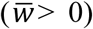 and 10% of 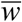 for inhibitory connections 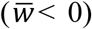. The average weights and probabilities of connections are specified in Table 2.

Heterogeneity of neurons within each population was provided by random distributions of the base values of the mean leakage reversal potentials *Ē*_Li0_ (see mean values ± SD for each *i*-th population in Table 1) and initial conditions for the values of membrane potential and channel kinetics variables. The base values of *Ē*_Li0_ and all initial conditions were assigned prior to simulations from their defined average values and variances using a random number generator, and a settling period of 10-200 s was allowed in each simulation.

#### 4.4.3 Simulations of changes in the locomotor frequency by neuroactive drugs and application of photostimulation

In the model, the frequency of rhythmic oscillations depended on the parameter α, that defined the level of average neuronal excitation in each population *i* (Shevtsova et al., 2015): *Ē*_Li_ = *Ē*_Li0_ × (1 − *α*) where *Ē*_Li0_ represents the base value of mean leakage reversal potential in the population at α=0 (see Table 1).

To simulate the effect of photostimulation, we selectively activated the V3 neurons either bi- or unilaterally by increasing the channelrhodopsin current conductance (*g*_chR_), which was set to 0 in control conditions. The values of *g*_chR_ for particular simulations are indicated in figure legends.

#### 4.4.4 Computer simulations

All simulations were performed using the custom neural simulation package NSM 4.1 (https://github.com/RybakLab/nsm) developed at Drexel University by S. N. Markin, I. A. Rybak and N. A. Shevtsova. This simulation package was previously used for the development of several spinal cord models (McCrea and Rybak, 2007, 2008; Rybak et al., 2006a, 2006b, 2013; Shevtsova et al., 2015; Shevtsova and Rybak, 2016; Zhong et al., 2012). Differential equations were solved using the exponential Euler integration method with a step size of 0.1 ms.

## 5 Author Contributions

SD, HZ, NS, IA, and YZ: conceptualization; SD, HZ, NS, IA, and YZ: methodology; YZ, HZ and JB: experiments; SD and NS: modeling; SD, HZ, and NS: formal analysis and software; HZ and SD: data curation; SD, HZ, NS, IA, and YZ: writing (original draft preparation); SD, HZ, NS, IA, and YZ: writing (review and editing); SD, HZ, and NS: visualization; IA and YZ: supervision, project administration, and funding acquisition.

## 6 Acknowledgements

This work was supported by CIHR grant MOP110950 and NSERC grant RGPIN 04880 (YZ) and NIH grants R01NS090919, R01NS095366, and R01NS100928 (IR).

